# COVID-19 adenoviral vector vaccination elicits a robust memory B cell response with the capacity to recognize Omicron BA.2 and BA.5 variants

**DOI:** 10.1101/2023.02.28.530547

**Authors:** Holly A. Fryer, Gemma E. Hartley, Emily S.J. Edwards, Nirupama Varese, Irene Boo, Scott J. Bornheimer, P. Mark Hogarth, Heidi E. Drummer, Robyn E. O’Hehir, Menno C. van Zelm

**Affiliations:** Allergy and Clinical Immunology Laboratory, Department of Immunology, Central Clinical School, Monash University, Melbourne, VIC, Australia; Immune Therapies Group, Burnet Institute, Melbourne, VIC, Australia; Viral Entry and Vaccines Group, Burnet Institute, Melbourne, VIC, Australia; BD Biosciences, San Jose, CA, USA; Department of Pathology, The University of Melbourne, Parkville, VIC, Australia; Department of Microbiology and Immunology, Peter Doherty Institute for Infection and Immunity, University of Melbourne, Melbourne, VIC, Australia; Department of Microbiology, Monash University, Clayton, VIC, Australia; Allergy, Asthma and Clinical Immunology Service, Alfred Hospital, Melbourne, VIC, Australia

**Author notes:** Corresponding author: Menno C. van Zelm, Department of Immunology, Central Clinical School, Monash University, 89 Commercial Road, Melbourne, VIC 3004, Australia.

## Abstract

Following the COVID-19 pandemic caused by SARS-CoV-2, novel vaccines have successfully reduced severe disease and death. Despite eliciting lower antibody responses, adenoviral vector vaccines are nearly as effective as mRNA vaccines. Therefore, protection against severe disease may be mediated by immune memory cells. We here evaluated plasma antibody and memory B cells (Bmem) targeting the Spike receptor binding domain (RBD) elicited by the adenoviral vector vaccine ChAdOx1 (AstraZeneca), their capacity to bind Omicron subvariants, and compared this to the response elicited by the mRNA vaccine BNT162b2 (Pfizer-BioNTech). Whole blood was sampled from 31 healthy adults pre-vaccination, and four weeks after dose one and dose two of ChAdOx1. Neutralizing antibodies (NAb) against SARS-CoV-2 were quantified at each timepoint. Recombinant RBDs of the Wuhan-Hu-1 (WH1), Delta, BA.2, and BA.5 variants were produced for ELISA-based quantification of plasma IgG and incorporated separately into fluorescent tetramers for flow cytometric identification of RBD-specific Bmem. NAb and RBD-specific IgG levels were over eight times lower following ChAdOx1 vaccination than BNT162b2. In ChAdOx1-vaccinated individuals, median plasma IgG recognition of BA.2 and BA.5 as a proportion of WH1-specific IgG was 26% and 17%, respectively. All donors generated resting RBD-specific Bmem, which were boosted after the second dose of ChAdOx1, and were similar in number to those produced by BNT162b2. The second dose of ChAdOx1 boosted Bmem that recognized VoC, and 37% and 39% of WH1-specific Bmem recognized BA.2 and BA.5, respectively. These data uncover mechanisms by which ChAdOx1 elicits immune memory to confer effective protection against severe COVID-19.

## Introduction

The coronavirus disease 2019 (COVID-19) pandemic is an ongoing worldwide health threat that has caused over 757 million cases and over 6.8 million deaths, as of February 2022.^1^ The virus causing COVID-19, severe acute respiratory syndrome coronavirus-2 (SARS-CoV-2), has a positive-sense single-stranded RNA genome which encodes four main structural proteins: Spike, Nucleocapsid (NCP), Envelope, and Membrane.^2^ Viral fusion is mediated by the receptor-binding domain (RBD) of the Spike protein, which binds the human angiotensin converting enzyme 2 (ACE2) receptor, initiating entry into host cells.^3, 4^

In efforts to control the pandemic there has been rapid global uptake of new mRNA and adenoviral vector vaccines, which are both over 90% effective against severe disease and hospitalization with COVID-19.^5, 6^ These COVID-19 vaccines use the Spike protein as the main immunogen, as the RBD is the major target for neutralizing antibodies (NAb).^7–11^ In Australia, the Pfizer-BioNTech BNT162b2 mRNA vaccine (BNT162b2), and the AstraZeneca ChAdOx1 nCoV-19 adenoviral vector vaccine (ChAdOx1) were first approved for two-dose primary schedules.^12, 13^ As evidence arose that protection against infection decreased over 4-6 months following two-dose vaccination, mRNA booster vaccines (3^rd^ and 4^th^ doses) were recommended.^14, 15^ Despite this, ChAdOx1 is the most widely distributed COVID-19 vaccine globally, meaning that a large proportion of the global population received this vaccine for their primary doses.^1, 16, 17^

It has been well-characterized that levels of NAb, Spike-specific and RBD-specific IgG detected after two doses of ChAdOx1 were all significantly lower than those elicited by mRNA vaccines such as BNT162b2.^18–21^ After either vaccine type, the RBD-specific antibody levels significantly declined after one-month post-vaccination; however, IgG remained detectable at least six months post-dose two.^22–24^ The Spike RBD-specific Bmem elicited by two doses BNT162b2 have been studied comprehensively.^22–27^ These Bmem are boosted by the second dose and show increased affinity of their surface Ig.^23^ Spike- and RBD-specific Bmem displayed a resting phenotype with the majority expressing surface CD27, and most lacking surface CD71, a marker of recent activation.^22, 26^ Spike-specific Bmem have also been detected in all individuals after two doses of ChAdOx1.^28^ However, it remains unclear how these Bmem compared to those elicited by mRNA vaccination with regards to their phenotype, durability, and kinetics.

Multiple SARS-CoV-2 variants of concern (VoC) have arisen, which evade the humoral immune response to differing degrees, mostly due to mutations in the RBD.^25, 29–33^ Following emergence of the Alpha, Beta, Gamma, and Delta variants, the currently designated VoC (as of January 2022) is the Omicron variant, with multiple sublineages including BA.1, BA.2, BA.4, and BA.5.^34–36^ Omicron demonstrates enhanced transmissibility and has been found to escape vaccine-elicited NAb to a far greater degree than any of the past VoC. The original Omicron sublineage BA.1 carries 35 Spike mutations, 15 of which are in the RBD.^37–40^ The BA.2 sublineage has 16 RBD mutations, with six differences from BA.1, and became the predominant SARS-CoV-2 variant around March 2022.^38, 39^ BA.4 and BA.5 have the same RBD sequence with 17 mutations and share all BA.2 RBD mutations except for Q493R, while having acquired L452R, shared with Delta, and F486V.^38, 39^ BA.5 overtook BA.2 as the prevalent circulating strain worldwide around August 2022.

Two doses of either an mRNA or adenoviral vector vaccine elicit very weak NAb against all Omicron sublineages, with levels 10-100-fold lower than those against the WH1 strain.^32, 41–52^ However, mRNA vaccines have been found to elicit substantial frequencies of Omicron-specific Bmem.^26, 27, 53^ Higher somatic hypermutation (SHM) levels were found in variant-binding Bmem than in those that only recognized the original WH1 virus, suggesting that continued maturation of high-affinity Bmem may improve recognition of highly mutated VoC.^22, 54^ The capacity of adenoviral vector vaccines to induce Bmem that bind these VoC is still unknown and is required to provide insight into whether the current primary vaccine schedules are sufficient to provide long-term protection against breakthrough infection with VoC. The global predominance of ChAdOx1 primary doses further emphasizes the importance of evaluating the durability of immune memory elicited by this vaccine.

Here, we evaluated the NAb, RBD-specific plasma IgG and circulating Bmem responses elicited by ChAdOx1 in a cohort of healthy adults (n=31) in the absence of SARS-CoV-2 infection using serological analyses and multi-parameter flow cytometry. Antibody and Bmem levels were quantified and compared to those elicited by BNT162b2. We also examined the ability of ChAdOx1-elicited IgG and Bmem to bind the RBDs from the Delta, Omicron BA.2, and BA.5 variants.

## MATERIALS AND METHODS

### Participants

From February to June 2021, 31 healthy adults without immunological or hematological disease or receiving immunomodulatory treatment and who received two doses of the ChAdOx1 vaccine were recruited to a low risk research study (**Supplementary Table 1**). Following written informed consent, 40mL of peripheral blood was collected pre-vaccination, four weeks post-dose one, and four weeks post-dose two of ChAdOx1 vaccination. Basic demographics including age, sex and COVID-19 infection status were collected throughout the study. This study was conducted according to the Declaration of Helsinki and approved by local human research ethics committees (Alfred Health ethics no. 32/21, Monash University project no. 72794).

### Protein production

DNA constructs encoding the SARS-CoV-2 RBD of WH1, Delta, and Omicron BA.2 and BA.5 were designed incorporating an N-terminal Fel d 1 leader sequence, a C-terminal AviTag for biotin ligase (BirA)-catalyzed biotinylation, and a 6-His tag for cobalt affinity column purification.^26, 55^ The DNA construct encoding the SARS-CoV-2 WH1 NCP protein was generated with an N-terminal human Ig leader sequence and the same C-terminal AviTag and 6-His tag.^55^

The DNA constructs were cloned into a pCR3 plasmid and produced using the Expi293 Expression system as described previously (Thermo Fisher, Waltham, MA).^26, 55^ While the NCP, WH1 RBD, and Delta RBD proteins were produced at 37°C,^26, 55^ production of the Omicron BA.2 and BA.5 RBD proteins was optimized at 34°C and with higher volume cultures. All proteins were purified, biotinylated and tetramerized with fluorochrome-conjugated streptavidin as described previously.^26, 55^ Briefly, biotinylated WH1 RBD was tetramerized by addition of either Brilliant Ultra Violet (BUV)395-conjugated streptavidin or BUV737-conjugated streptavidin, and biotinylated Omicron BA.2 and BA.5 were tetramerized with Brilliant Violet (BV)480-conjugated streptavidin and BV650-conjugated streptavidin respectively, at a protein:streptavidin molar ratio of 4:1 to produce [RBD WH1]_4_-BUV395, [RBD WH1]_4_-BUV737, [RBD BA.2]_4_-BV480 and [RBD BA.5]_4_-BV650 (**Supplementary Tables 2 and 3**).^26, 55^

### Pseudovirus neutralization assay

Plasma NAb levels against the WH1 virus and VoC (Delta and Omicron BA.2 and BA.5) were evaluated using SARS-CoV-2 retroviral pseudotyped viruses and a 293T-ACE2 cell line as previously described.^55^ The percentage entry was calculated as described previously and plotted against reciprocal plasma dilution using GraphPad Prism 9 Software (GraphPad Software, La Jolla, CA) and curves fitted with a one-site specific binding Hill plot.^55^ The reciprocal dilution of plasma required to prevent 50% virus entry was calculated from the non-linear regression line (IC50). The lowest detectable NAb titer was 20, and all samples that did not achieve 50% neutralization at this dilution were reported as the arbitrary value of 10.

### IgG ELISA

ELISAs were performed for each plasma sample to measure plasma IgG levels specific for the WH1 NCP and RBD and the VoC RBDs. 96-well EIA/RIA plates (Corning Incorporated, Costar, St Louis, MO) were coated with 2μg/mL unbiotinylated monomer RBD or NCP protein overnight at 4°C. Plates were then blocked with 3% BSA in PBS. Plasma was diluted 1:30 for quantification of RBD- and NCP-specific antibodies pre-vaccination, post-dose one and post-dose two. Plasma was titrated from 1:30 to 1:10,000 for quantification of WH1

RBD-specific antibodies post-dose one and WH1 and VoC (Delta and Omicron BA.2 and BA.5) RBD-specific antibodies post-dose two. Serially diluted human IgG (in-house made rituximab) was added to wells on the same plate to create a standard curve for quantification of IgG levels. Titration curves were created and the area under the curve (AUC) values for each variant were calculated using GraphPad Prism 9. IgG binding to VoC RBD was quantified relative to WH1 as a percentage of WH1 AUC.

### Flow cytometry

#### Trucount

Blood samples were processed as previously described.^55^ Absolute numbers of major leukocyte populations in each sample were determined as previously described.^26, 55, 56^ Trucount data were used to calculate the absolute numbers of RBD-specific Bmem subsets.^56^ Briefly, 50μL of whole blood was added to a BD Trucount tube (BD Biosciences, San Jose, CA, USA) and incubated with 20μL of the Multitest^TM^ 6-color TBNK reagent (BD Biosciences) containing CD3, CD4, CD8, CD19, CD16, CD45 and CD56 antibodies (**Supplementary Tables 2 and 3**) for 15 minutes in the dark at room temperature. Subsequently, cells were incubated with 1X BD Lysis Solution (BD Biosciences) for 15 minutes to lyse red blood cells. Samples were acquired on the BD FACSLyric analyzer and data was analyzed using FlowJo^TM^ Software v10.8.1 (BD Biosciences) as previously described.^55, 56^

#### RBD-specific Bmem analysis

Fluorescent tetramers of WH1, Omicron BA.2 and BA.5 RBDs were used to detect antigen-specific Bmem. 10-15×10^6^ thawed PBMC were incubated at room temperature in the dark for 15 minutes in a total volume of 250μL with fixable viability stain 700, antibodies against CD3, CD19, CD38, CD27, CD21, CD71, IgG1, IgG2, IgG3, IgG4, IgD, and IgA, and 5μg/mL each of [RBD WH1]4-BUV395, [RBD WH1]4-BUV737, [RBD BA.2]4-BV480, and [RBD BA.5]_4_-BV650, and FACS buffer (**Supplementary Tables 2 and 3**). In a separate tube, 1-5×10^6^ PBMCs were incubated with cocktail containing fixable viability stain 700, antibodies against CD3, CD19, CD27, and IgD, and fluorochrome-conjugated streptavidin controls (BUV395, BUV737, BV480, and BV650) without the RBD. Cells were then washed with FACS buffer, fixed with 2% PFA for 20 minutes at room temperature in the dark, washed once more and acquired on the LSRFortessa X-20 (BD Biosciences). Flow cytometry setup was performed using standardized EuroFlow settings as previously published, to ensure comparability of data.^26, 55–57^ Data analysis was performed using FlowJo^TM^ Software v10.8.1 (BD Biosciences).

### Statistical analysis

The absolute numbers of each B cell subpopulation were calculated relative to the total B cell count from the Trucount analysis. Statistical tests were performed using GraphPad Prism 9 Software. Paired data were analyzed using the non-parametric Wilcoxon signed ranks test for matched pairs. The non-parametric Mann-Whitney test was used to determine significance of unpaired data, and Spearman’s rank correlation was used to determine the correlation coefficient (R_S_) between two variables. For all tests, p<0.05 was considered significant.

## RESULTS

### Cohort characteristics

Peripheral blood was collected from 31 healthy adults (median age 45 years, range: 26-65 years; 74% female) before and after two doses of the ChAdOx1 vaccine, received with a median of 84 days between doses (range: 70-95 days) (**Figure 1A, Supplementary Table 1**). Pre-vaccination samples were taken in March and April 2021 from 30 donors, and post-vaccination samples were obtained from all 31 donors 28 days after dose 1 (range, 25-29 days) and 28 days after dose 2 (range, 27-36 days).

**Figure 1.**
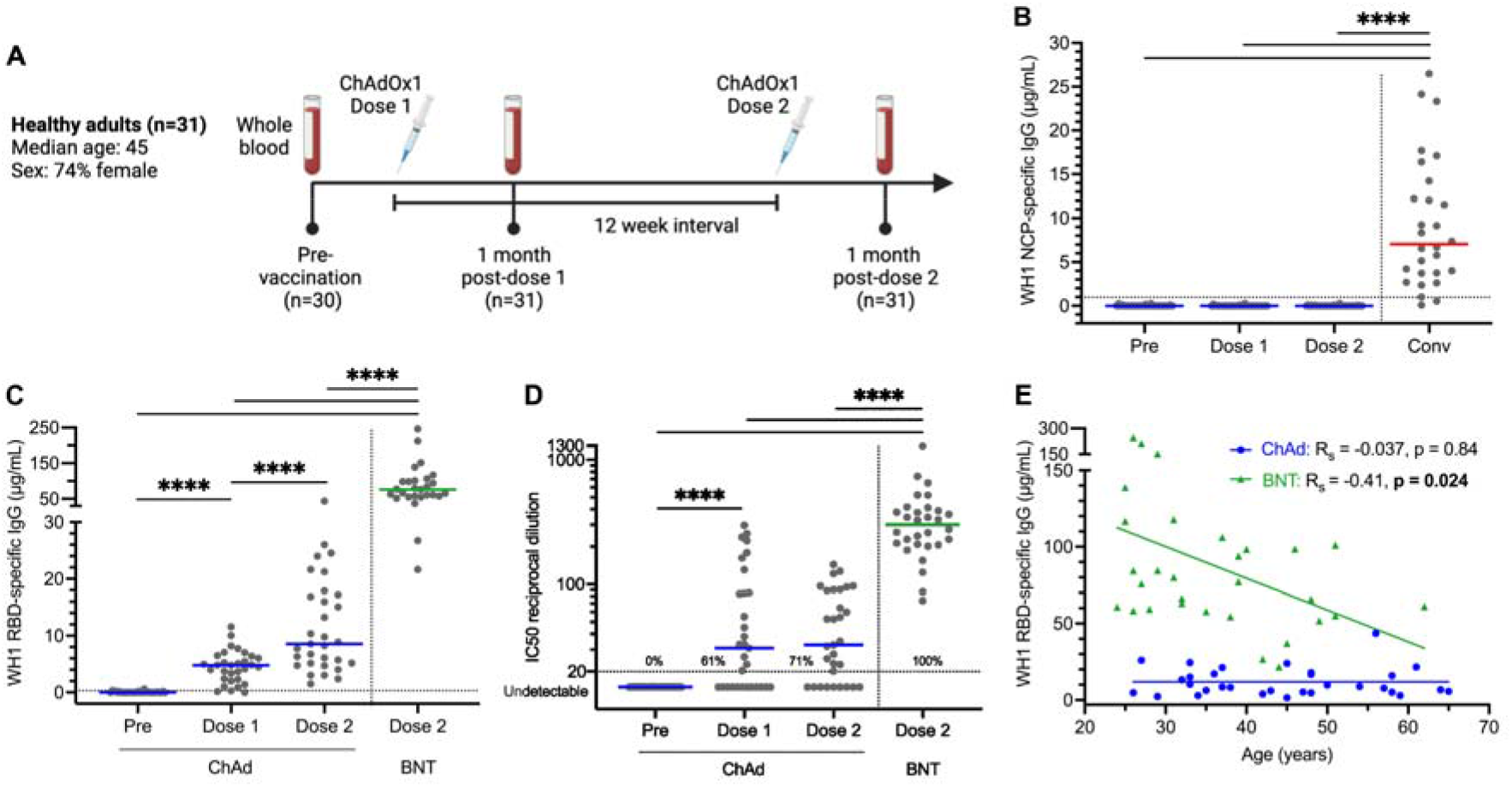
SARS-CoV-2 WH1-specific serological responses elicited by ChAdOx1 and comparison to BNT162b2. (**A**) Scheme of study design involving longitudinal sampling of 31 ChAdOx1 recipients pre-vaccination, four weeks post-dose one (median 28 days, range: 25-29), and four weeks post-dose two (median 28 days, range: 27-36 days), with a median of 84 days between doses (range: 70-95 days). Created in BioRender.com (2022). (**B**) WH1 NCP-specific and (**C**) RBD-specific plasma IgG concentrations pre-vaccination (Pre), four weeks post-dose one (Dose 1) and four weeks post-dose two (Dose 2) of ChAdOx1 (ChAd). BNT162b2 (BNT) post-dose two data added for statistical comparison (n=30).^26^ Convalescent (Conv) data previously published, added to illustrate NCP-specific response following SARS-CoV-2 infection (n=28).^55^ Horizontal dotted lines at 0.97μg/mL in (**B**) and 0.31μg/mL in (**C**) indicate the 10^th^ percentile of previously published SARS-CoV-2-convalescent patients.^55^ (**D**) NAb against WH1 SARS-CoV-2 generated by ChAdOx1. Horizontal dotted line indicates the neutralization cut-off at an IC50 value of 20.^55^ Values indicate the percent of donors producing neutralizing levels of antibody. Solid lines indicate median values. Mann-Whitney test for unpaired data and Wilcoxon matched-pairs signed rank test for paired data. Only significant differences shown. ****p<0.0001. (**E**) Correlation of WH1 RBD-specific IgG with age after either ChAdOx1 (n=31) or BNT162b2 (n=30) two-dose vaccination. Non-parametric Spearman’s rank correlation (Rs), solid line represents simple linear regression line. Convalescent and BNT162b2 data previously published.^26, 55^

### Double-dose ChAdOx1 elicits low levels of RBD-specific IgG and NAb

All 31 donors self-reported to be SARS-CoV-2 naive before and for the duration of the study. To independently confirm this, serology was performed for NCP, a SARS-CoV-2 antigen that is not present in the ChAdOx1 vaccine, therefore indicating previous SARS-CoV-2 infection. All donors were negative for NCP-specific IgG pre-vaccination, at four weeks post-dose one, and at four weeks post-dose two, with significantly lower levels than COVID-19 convalescent patients (**Figure 1B**).^55^ Thus, our cohort was proven to be SARS-CoV-2 infection naive.

Plasma IgG to WH1 RBD was undetectable in all donors before vaccination. After one dose, the concentration of WH1 RBD-specific IgG increased in 30/31 donors (97%), with a positive IgG response detected in 28/31 donors (90%) (**Figure 1C**). After the second dose, levels increased in 28/31 donors (90%), and a positive RBD-specific IgG response was detected in all donors (**Figure 1C**). Overall, WH1 RBD-specific IgG concentrations were significantly higher after dose two than pre-vaccination or after dose one (**Figure 1C**). However, the median WH1 RBD-specific IgG level of 8.6μg/ml after two doses of ChAdOx1 was almost nine-fold lower than that elicited by two doses of BNT162b2 (75.9μg/ml).^26^ Neutralizing antibodies (NAb) against the WH1 strain, as determined with a pseudovirus neutralization assay,^55^ were detectable in 19/31 donors (61%) post-dose one, and increased to 22/31 donors (71%) post-dose two (**Figure 1D**). In 5/31 ChAdOx1 recipients (16%), NAb levels were below the neutralization cutoff at all timepoints. In contrast, all BNT162b2 recipients (30/30) produced NAb against the WH1 strain after two doses, and BNT162b2 also elicited significantly higher NAb than ChAdOx1 (**Figure 1D**).

As the median age of the ChAdOx1 recipient cohort (45 years) was higher than that of the BNT162b2 cohort (32 years), correlation analysis was performed to determine whether age was a contributing factor to the lower serological response in ChAdOx1 recipients. WH1 RBD-specific IgG levels were consistently lower in ChAdOx1 recipients, irrespective of their age (**Figure 1E**). Remarkably, there was a slight negative correlation between WH1 RBD-specific IgG and age in the BNT162b2 recipients (p=0.024), whereas there was no correlation in the ChAdOx1 group (p=0.84).

### ChAdOx1 vaccination elicits similar numbers of WH1 RBD-specific Bmem to BNT162b2

To evaluate the capacity of ChAdOx1 vaccination to elicit SARS-CoV-2 specific Bmem, PBMC samples from all donors after both doses were extensively immunophenotyped with a 16-color flow cytometry panel including B-cell markers (CD27, CD21, CD38 and CD71), Ig isotype and IgG subclass antibodies (IgD, IgA, IgG1, IgG2, IgG3 and IgG4), and fluorescently-tagged RBD tetramers (**Supplementary Tables 2 and 3**). The panel facilitated enumeration of naive B cells (CD27^-^IgD^-^), unswitched (CD27^+^IgD^+^) and Ig class switched (CD27^+/-^IgD^-^) Bmem subsets (**Supplementary Figure 1**). RBD-specific B cells were identified by double-discrimination using both [RBD WH1]_4_-BUV395 and [RBD WH1]_4_-BUV737 tetramers to reduce background by excluding cells non-specifically binding a single fluorochrome. RBD-specific B cells were then categorized into subsets using the gating strategy as for total B cells (**Figure 2A**).

**Figure 2.**
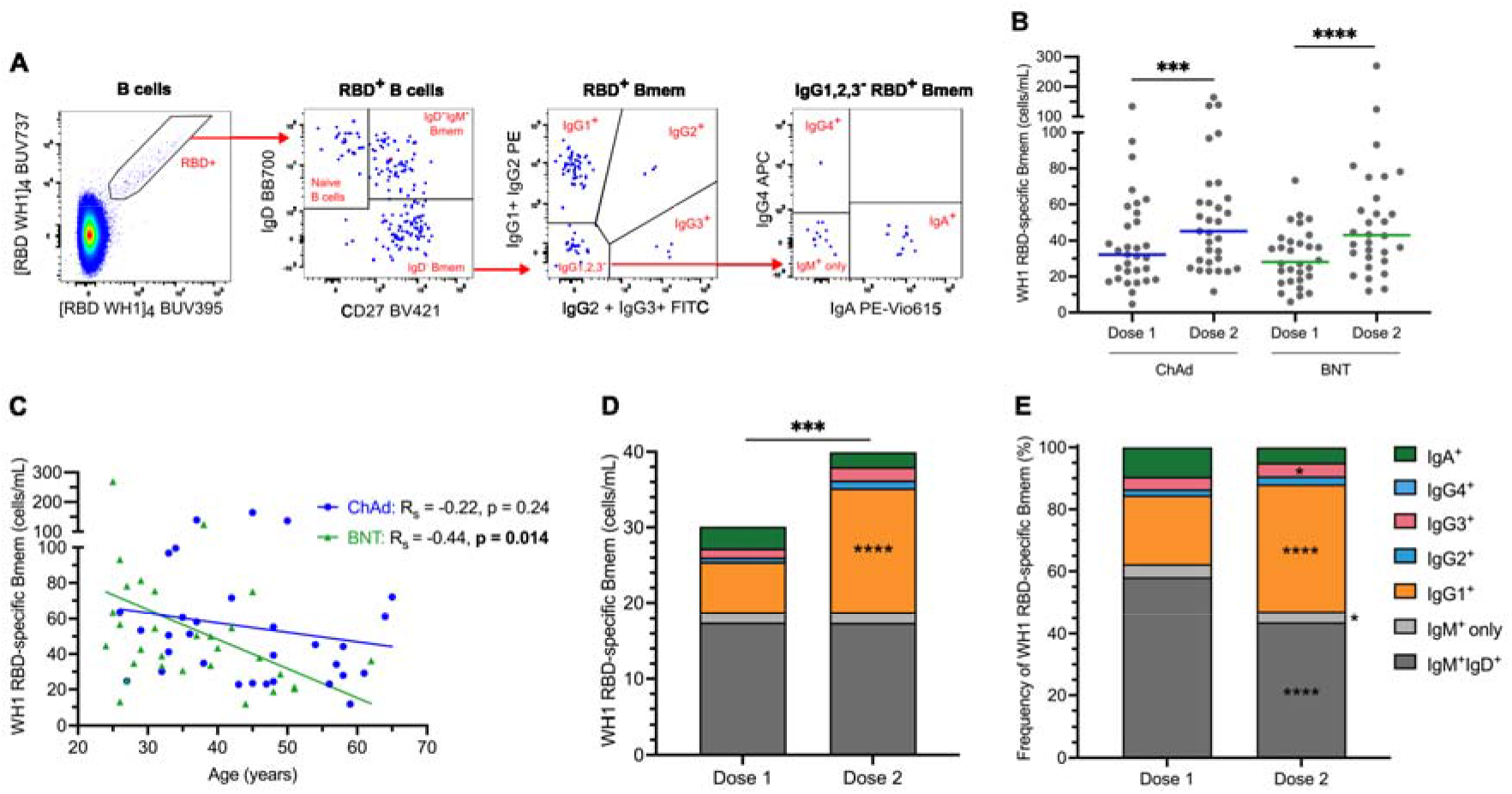
Robust RBD-specific Bmem response elicited by ChAdOx1 with an expansion of IgG1^+^ Bmem after dose 2. (**A**) Double-discrimination of WH1 RBD-specific B cells was achieved by gating B cells double-positive for [RBD WH1]_4_-BUV395 and [RBD WH1]_4_-BUV737. WH1 RBD-specific B cells were gated for CD27^-^IgD^+^ naive B cells, CD27^+^IgD^+^IgM^+^ natural effector cells, and IgD^-^ Bmem, then each subset was gated for IgG1^+^, IgG2^+^, IgG3^+^, IgG4^+^, IgA^+^, and IgM^+^ only cells. (**B**) Absolute numbers of WH1 RBD-specific Bmem four weeks post-dose one (Dose 1) and four weeks post-dose two (Dose 2) of ChAdOx1 (ChAd) and BNT162b2 (BNT). (**C**) Correlation of WH1 RBD-specific Bmem with age after either ChAdOx1 or BNT162b2 two-dose vaccination. Non-parametric Spearman’s rank correlation (Rs), solid line represents simple linear regression line. (**D-E**) The absolute numbers (**D**) and frequencies (**E**) of WH1 RBD-specific Bmem expressing IgM only, IgM and IgD, IgG1, IgG2, IgG3, IgG4, or IgA postvaccination with ChAdOx1. ChAdOx1 n=31, BNT162b2 n=30. Mann-Whitney test for unpaired data and Wilcoxon matched-pairs signed rank test for paired data. Only significant differences shown. *p<0.05, ***p<0.001, ****p<0.0001. BNT162b2 data previously published.^26^

Total RBD-specific Bmem were defined following exclusion of naive (IgD^+^CD27^-^) B cells (**Figure 2A**). RBD-specific Bmem were detected in all donors after dose one of ChAdOx1, and these numbers significantly increased after dose two (**Figure 2B**). In contrast to plasma IgG levels, there were no differences in median numbers of RBD-specific Bmem between ChAdOx1 and BNT162b2 vaccinees both after dose one and after dose two.

Correlation analysis was performed to determine the effect of donor age on the RBD-specific Bmem levels of the two vaccine groups after dose two (**Figure 2C**). There was no significant correlation between Bmem numbers with age in the ChAdOx1 group (p=0.24), while in the BNT162b2 group there was a slight, but significant (p=0.014) negative correlation. Therefore, the slightly higher median age of the ChAdOx1 cohort did not affect the comparison of Bmem cells to the BNT162b2 cohort.

Detailed immunophenotype analysis of the RBD-specific Bmem compartment revealed that the expansion after dose two of ChAdOx1 was solely due to increased numbers of IgG1^+^ Bmem cells (**Figure 2D**). The absolute number of unswitched RBD-specific Bmem, as well as those expressing IgG2, IgG3, IgG4 or IgA, were similar after both doses. The expansion of the frequency of IgG1^+^ RBD-specific Bmem between the two timepoints was accompanied by a relative decrease in the proportions of IgM^+^IgD^+^, IgM^+^ only, and IgA^+^ RBD-specific Bmem, while the absolute numbers of these populations were similar (**Figure 2E**). Neither age nor individual donors appeared to affect the expansion of IgG1^+^ WH1 RBD-specific Bmem, as the IgG1^+^ population increased in number in 29/31 individuals (94%) after the second dose of ChAdOx1 (**Supplementary Figure 2**). Additionally, there was a weak positive correlation between RBD-specific serum IgG levels and IgG^+^ RBD-specific Bmem number after one dose of ChAdOx1 (R_s_=0.45, p=0.012), and when data from both doses were combined (R_s_=0.43, p=0.0005), but not after dose two alone (**Supplementary Figure 3**). Despite the significant increase in RBD-specific Bmem numbers, total Bmem numbers, as well as their composition based on Ig isotype and IgG subclass, were similar after dose one and dose two of ChAdOx1 (**Supplementary Figure 4A-D**). This demonstrates that the phenotype of the total Bmem compartment was not affected by the vaccination, and that ChAdOx1 vaccination induced the expansion of only RBD-specific IgG1^+^ Bmem.

### The ChAdOx1 vaccine-elicited Bmem display a resting memory phenotype

The phenotype of total and RBD-specific Bmem following ChAdOx1 vaccination was characterized using a range of memory and activation markers (**Figure 3A-C**). Expression of CD27 on IgG^+^ B cells marks continued maturation in the germinal center (GC), linked to higher SHM levels than in CD27^-^ Bmem.^58^ The median frequency of IgG^+^ RBD-specific Bmem expressing CD27 significantly increased from 81% four weeks after dose one to 88% four weeks after dose two of ChAdOx1 (**Figure 3D**). Low expression of CD21 on B cells is associated with recent activation, distinct from classical, quiescent CD21^+^ Bmem.^59, 60^ The median frequencies of CI)21^lo^ cells within RBD-specific Bmem remained low at 9% at both timepoints, with no significant change after dose two compared to dose one (**Figure 3E**). Expression of CD71 marks recently activated B cells.^59^ Median frequencies of CD38^dim^CD71^+^ cells within RBD-specific Bmem significantly decreased from 2% after dose one to 1% after dose two of ChAdOx1 (**Figure 3F**). Within total Bmem, there were no significant changes in the frequencies of CD27^+^IgG^+^ Bmem, CD21^lo^ Bmem, or CD38^dim^CD71^+^ Bmem between post-dose one and post-dose two samples, representing the stability of the total Bmem pool (**Supplementary Figure 4E-G**). Overall, this indicates that one month after two doses of ChAdOx1 a mature, quiescent SARS-CoV-2-specific Bmem population was formed.

**Figure 3.**
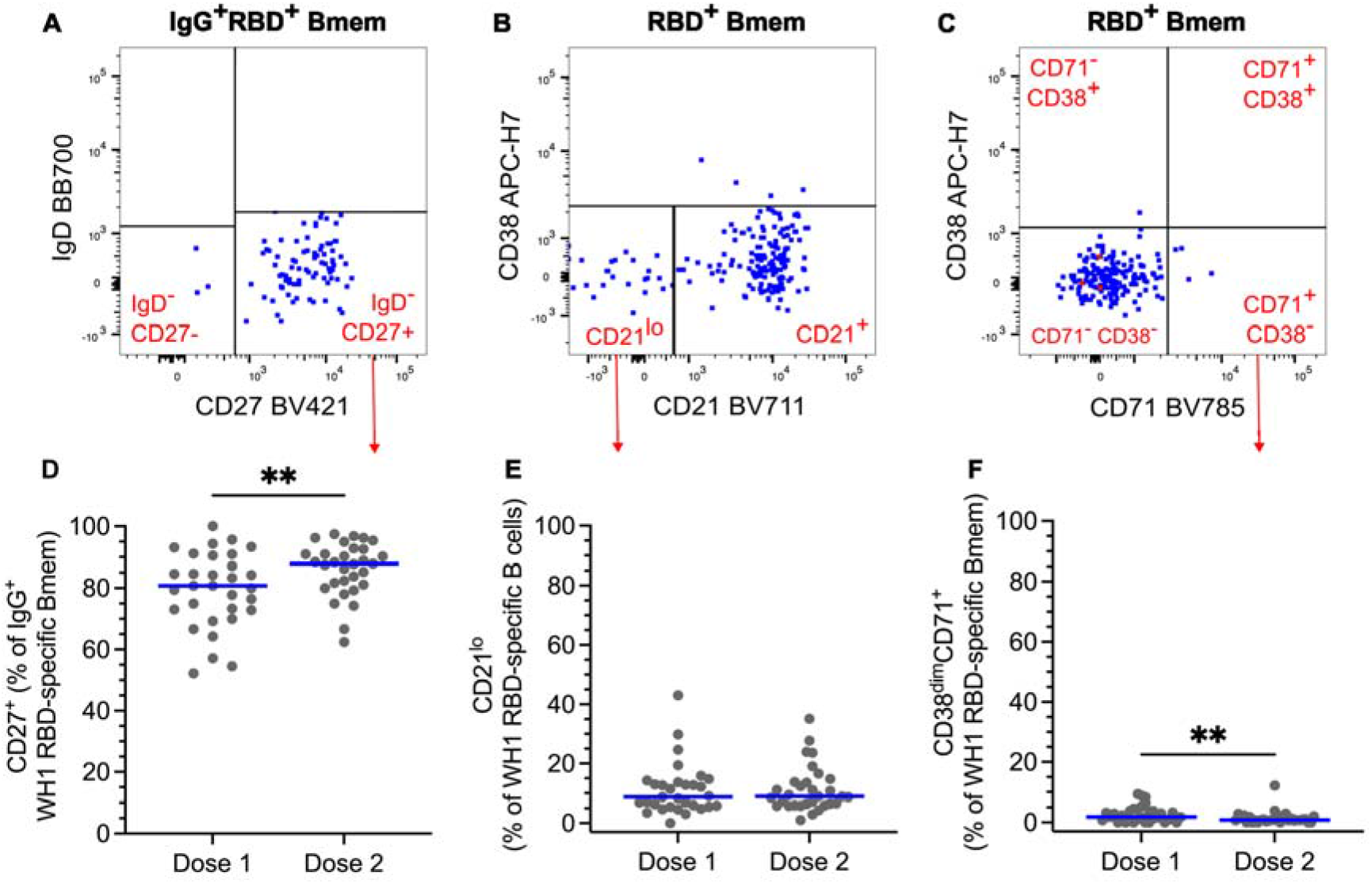
WH1 RBD-specific Bmem with a resting phenotype are elicited by two doses of ChAdOx1 vaccination. (**A**) IgG^+^ WH1 RBD-specific Bmem were gated for the expression of CD27. WH1 RBD-specific Bmem were gated for (**B**) CD21 and (**C**) CD71 expression. (**D**) Frequencies of IgG^+^ WH1 RBD-specific Bmem expressing CD27 four weeks post-dose one (Dose 1) and four weeks post-dose two (Dose 2) of ChAdOx1. (**E-F**) The frequencies of (**E**) CD21^lo^ and (**F**) CD38^dim^CD71^+^ WH1 RBD-specific Bmem post-vaccination with ChAdOx1. n=31. Wilcoxon matched-pairs signed rank test. Only significant differences shown. **p<0.01.

### The second dose of ChAdOx1 increases the capacity of RBD-specific Bmem to recognize Omicron BA.2 and BA.5

After two doses of ChAdOx1, all donors had detectable plasma IgG that bound WH1 RBD, therefore we next evaluated the capacity of this antibody response to recognize Delta, Omicron BA.2, and BA.5 VoC. While there was no difference in the levels of IgG specific for the Delta and WH1 RBDs, the levels of IgG specific for BA.2 or BA.5 were significantly decreased compared to WH1 (**Figure 4A**). The median proportions of WH1-specific IgG that bound Delta, BA.2, and BA.5 were 90%, 26%, and 17%, respectively (**Figure 4A**). Levels of WH1-specific IgG that recognized either Omicron sublineage were significantly lower than Delta-specific IgG, and IgG that bound BA.5 was significantly lower than that which bound BA.2 (**Figure 4A**).

**Figure 4.**
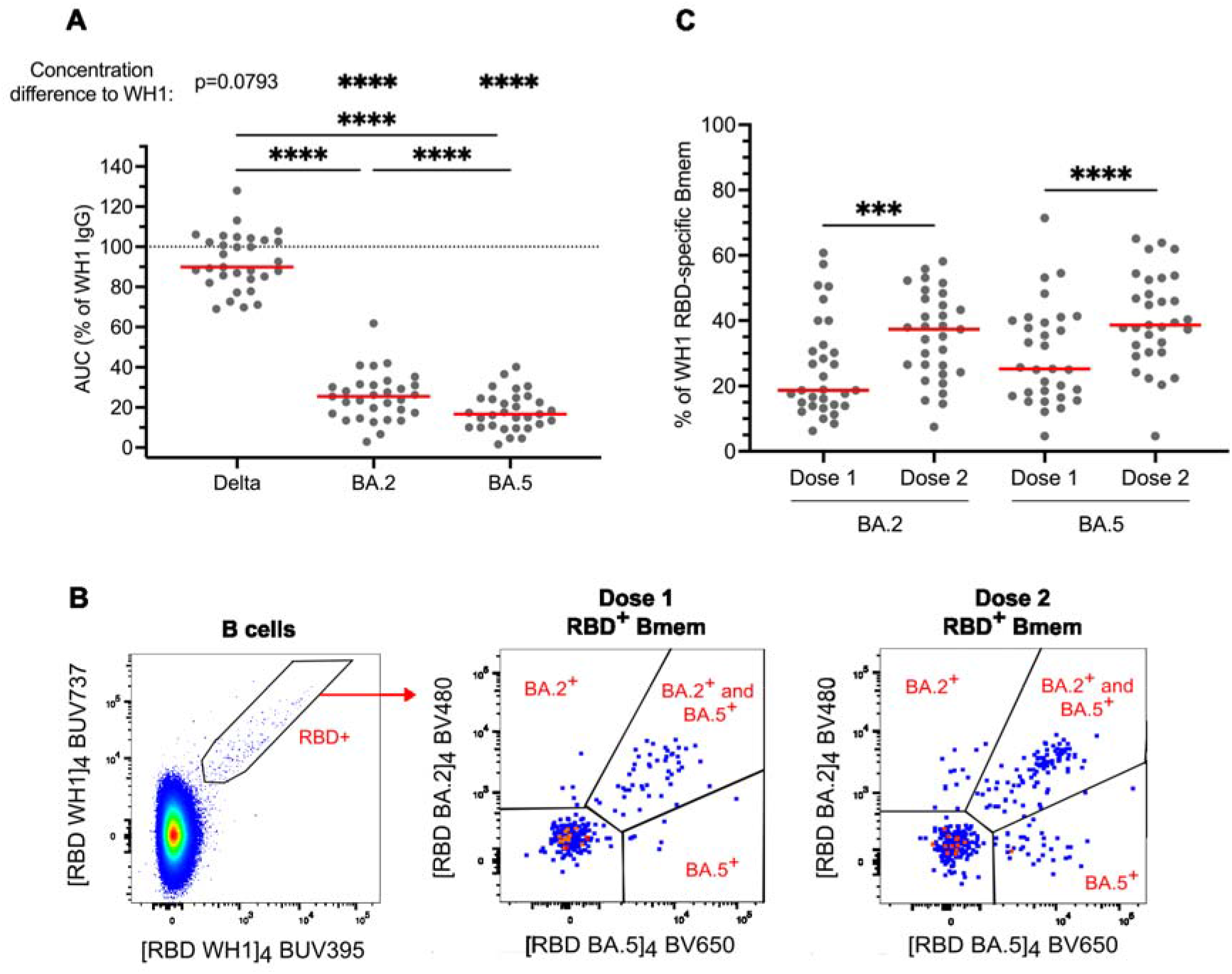
Two doses of ChAdOx1 elicit plasma IgG and RBD-specific Bmem to recognize variants of concern. (**A**) Plasma IgG specific for Delta, BA.2 and BA.5 RBD expressed as a percentage of WH1-specific IgG post-dose two of ChAdOx1. Dotted line indicates 100% of WH1 IgG. (**B**) Gating strategy to enumerate WH1 RBD-specific Bmem that bound BA.2 and/or BA.5 RBD four weeks post-dose one and four weeks post-dose two of ChAdOx1. (**C**) Frequencies of WH1 RBD-specific Bmem that bound BA.2 and/or BA.5 RBD post-vaccination with ChAdOx1 (n=31). Solid lines indicate median values. Wilcoxon matched-pairs signed rank test. Only significant differences shown. ***p<0.001, ****p<0.0001.

To evaluate the capacity of Bmem cells to recognize VoC, detection of cells binding BA.2 and BA.5 RBD tetramers was evaluated within the WH1 RBD-specific Bmem population by flow cytometry four weeks after dose one and four weeks after dose two of ChAdOx1 (**Figure 4B**). After dose one, the median percentage of WH1 RBD-specific Bmem that bound BA.2 was 19%, and the percentage that bound BA.5 was 25% (**Figure 4C**). The second vaccine dose significantly boosted these frequencies, increasing the percentage of WH1 RBD-specific Bmem numbers binding BA.2 to 37%, and those binding BA.5 to 39%. This increase in the proportion of variant-binding RBD-specific Bmem was mainly due to a significant increase in IgG1^+^ RBD-specific Bmem that bound BA.2 or BA.5 (**Supplementary Figure 5**). The majority of variant-binding Bmem recognized both BA.2 and BA.5 fell (i.e double-positive), 63% after dose one and 69% after dose two (data not shown). In comparison to BNT162b2, ChAdOx1 elicited similar numbers of BA.2-binding RBD-specific Bmem at both timepoints, but higher numbers of BA.5-binding RBD-specific Bmem after dose two (**Supplementary Figure 6**). Overall, after two doses of ChAdOx1, significantly higher proportions of RBD-specific Bmem bound BA.2 (37%) and BA.5 (39%) than RBD-specific plasma IgG did (19 and 25% respectively). This indicates that the RBD-specific Bmem compartment has a greater capacity to recognize highly-mutated VoC than the circulating RBD-specific plasma IgG.

## DISCUSSION

We here present the RBD-specific plasma IgG and Bmem responses in healthy SARS-CoV-2 naive adults who received two doses of the adenoviral vector COVID-19 ChAdOx1 vaccine. We have demonstrated that whilst the antibody response is suboptimal, the two-dose primary schedule of ChAdOx1 induces a robust, resting RBD-specific Bmem population, which is not significantly affected by age and has the capacity to recognize highly mutated VoC. The magnitude and variant-binding capacity of the RBD-specific Bmem population was boosted by the second dose of ChAdOx1. RBD-specific Bmem numbers and their capacity to bind Omicron BA.2 did not differ from mRNA vaccine BNT162b2 recipients, and the capacity of these Bmem to bind BA.5 was greater in ChAdOx1 recipients.

After the second dose of ChAdOx1, all our 31 donors produced detectable WH1 RBD-specific IgG, and the median WH1 RBD-specific IgG levels approximately doubled. These trends are in line with previous studies reporting that all recipients of double-dose ChAdOx1 seroconvert.^18, 19, 61, 62^ ChAdOx1 only elicited detectable NAb in 71% of donors after two doses, which is contradicted by the original phase one study and two subsequent clinical trials that found 99-100% of double-dose ChAdOx1 recipients produced detectable NAb.^63, 64^ However, the clinical trials utilized live SARS-CoV-2 assays whereas we performed a pseudovirus assay, and each assay defines different neutralization cut-offs, making it difficult to directly compare NAb titers. In addition, some trials measured NAb levels only two weeks after the second dose, whereas we sampled four weeks after, when antibody levels have been shown to begin waning.^65, 66^

ChAdOx1 recipients generated significantly lower WH1 RBD-specific plasma IgG and NAb levels than BNT162b2 recipients after dose two, in line with the current literature.^18, 19, 61^ One mechanism behind this may be the difference in Spike protein structures; the mRNA in BNT162b2 encodes the prefusion form, stabilized by two proline substitutions, while the DNA in ChAdOx1 encodes the wild-type, unmodified protein.^67, 68^ The stabilizing mutations in BNT162b2 prevent the shedding of the S1 subunit from the Spike protein, and this conformation has been shown to be more immunogenic.^6, 69–71^ Another factor that can influence the antibody response elicited by vaccines is anti-vector immunity, such as the generation of NAb specific for the adenovirus backbone component of ChAdOx1. This response can inhibit the amount of vaccine antigen that is delivered, resulting in lower immunogenicity.^72, 73^ It is unlikely that this is the driving mechanism in the case of ChAdOx1, as the Y25 adenovirus in the vaccine is chimpanzee-derived and thus has low seroprevalence in humans.^74^ However, it is possible that additional third or fourth adenoviral vector vaccine doses may augment this effect and induce higher levels of vector-specific antibodies, inhibiting the response to the Spike protein. This would suggest that it may be more effective to use a different formulation such as mRNA vaccines for subsequent doses, which is already recommended in many countries due to the link between ChAdOx1 and events of thrombosis.^75^

The diminished serological response elicited by ChAdOx1 compared to BNT162b2 may be related to different vaccine efficacies. While the phase three trials were not performed head to head, the efficacy of ChAdOx1 against symptomatic infection after two doses was 70.4% compared to 95% for BNT162b2.^6, 76^ However, the two vaccines have similarly high efficacy rates against severe disease and hospitalization with COVID-19, at 100% after two doses of ChAdOx1 and 92% after BNT162b2.^76–78^ We found that the two vaccines elicited similar numbers of WH1 RBD-specific Bmem after each dose, suggesting that Bmem may be a better indicator of long-term protection against severe COVID-19 than antibody levels. Interestingly, we found that there was some correlation between IgG^+^ WH1 RBD-specific Bmem and WH1 RBD-specific IgG levels following ChAdOx1 vaccination. However, it was not as strong as the correlation after BNT162b2 vaccination that we reported previously.^26^

Through the measurement of absolute cell counts, we found no difference between the numbers of RBD-specific Bmem elicited by ChAdOx1 and BNT162b2 one month post-dose two. In contrast to these results, Wang *et al*. (2022) performed a similar assay following two-dose vaccination, sampling five months after the second dose, and found ChAdOx1 elicited lower frequencies of RBD-specific B cells than BNT162b2.^66^ Other studies have also reported lower frequencies of Spike- or RBD-specific Bmem following adenoviral vector COVID-19 vaccination compared to mRNA vaccination.^70, 79^ However, frequencies can be impacted by changes in the numbers of other cell populations, making the measurement less robust than absolute numbers. Additionally, our current study measured a post-dose two timepoint four months earlier than Wang *et al*. (2022).^66^ Together, this makes it difficult to directly compare these findings with our own. The persistence of WH1 RBD-specific Bmem following ChAdOx1 vaccination is still unknown and would be a relevant subject of further studies to determine the durability of vaccine-induced immune memory.

The 12-week interval between the first and second ChAdOx1 vaccine dose was much longer than the three weeks between BNT162b2 doses. Delaying the second mRNA vaccine dose has been shown to increase circulating NAb and RBD-specific antibody. Additionally, a longer dosing interval may augment the Bmem pool generated by the first dose, as reduced antigen availability over time caused by a delayed second dose may increase competition in the GC and improve Bmem quality through affinity maturation.^80^ This suggests that while the antibody response elicited by ChAdOx1 is lower after two doses, the enhanced Bmem pool which matured during the extended dosing interval could improve antibody production upon the receipt of a third dose. This is supported by the literature, as recipients of two doses of either vaccine type have been shown to produce similar NAb titers after receiving an mRNA booster dose.^81^ Comparable Bmem numbers between the two cohorts after two doses would translate into similar NAb levels after the third dose, as cells within the resting Bmem pool are reactivated and induced to differentiate into antibody-secreting cells. Thus, both dosing interval length and vaccine formulation may impact the magnitude of Bmem and humoral immune responses.

The RBD-specific Bmem elicited by ChAdOx1 exhibited a similar resting phenotype as following BNT162b2 vaccination, with low CD71 expression and high expression of CD27 within IgG^+^ RBD-specific Bmem.^26^ This indicates that both vaccines have the capacity to elicit an antigen-specific Bmem population that is predominantly resting and Ig class switched by one-month post-dose two, likely having undergone further maturation in a GC after reactivation. Thus, these RBD-specific Bmem are primed for a robust recall response. After the second dose of ChAdOx1, there was a significant increase in the absolute number of IgG1^+^ RBD-specific Bmem, while Bmem of all other isotypes and IgG subclasses remained unchanged in number, similar to the response elicited by BNT162b2.^26^ This is indicative of a T-dependent B cell response to viral protein antigen, which typically skews the IgG subclass distribution toward potent viral neutralizers IgG1 and IgG3^82–85^. These were the two most prevalent IgG subclasses of RBD-specific Bmem following ChAdOx1 vaccination, demonstrating that the vaccine stimulates a Bmem population with the capacity for producing potent NAb upon differentiation.

After two doses of ChAdOx1, IgG and Bmem recognition of Omicron BA.2 and BA.5 VoC RBDs were reduced compared to WH1, illustrating escape from the vaccine-induced immune response caused by RBD mutations. Additionally, the percentage of circulating IgG that bound BA.5 was lower than BA.2, indicating a higher degree of escape from antibody binding by BA.5, which is in line with recent reports.^47–50, 52^ In particular, RBD mutations including L452R and F486V have been shown to increase the ability of BA.5 to escape from serum neutralization.^33, 48^ This helps to explain the global predominance of the sublineage in late 2022 and its capacity for rapid transmission even in largely vaccinated populations.^48, 50, 52^ Following ChAdOx1 vaccination, the levels of RBD-specific IgG that bound both BA.2 and BA.5, relative to WH1-specific IgG, were similar compared to those elicited by BNT162b2.^26^ ChAdOx1 also elicited similar frequencies of BA.2-specific Bmem to BNT162b2 after both the first and second doses, but higher frequencies of BA.5-specific Bmem after dose two.^26^ This indicates that both the mRNA and adenoviral vector vaccine formulations can elicit an immune memory response capable of recognizing current VoC.

The frequencies of RBD-specific Bmem that recognized BA.2 and BA.5 after two vaccine doses were higher than the frequencies of RBD-specific plasma IgG that bound each VoC. One functional benefit that Bmem possess over terminally differentiated plasma cells, which have fixed specificity, is the capacity to evolve their antibody binding affinity and breadth. This occurs at each subsequent antigen encounter, either a booster vaccination or reinfection, when GC reactions lead to increased SHM in Bmem.^86^ These processes could explain why ChAdOx1 elicits a resting Bmem population with an increased capacity to bind highly-mutated viral variants compared to the circulating IgG pool.^87^

The discrepancy between the median ages of the ChAdOx1 and BNT162b2 recipients compared in this study (45 and 32 years, respectively) was largely unavoidable, as it was more likely that older individuals would receive ChAdOx1 due to Australian Government recommendations.^88^ As the immune response can decline with age and others have shown that Spike-specific antibodies were diminished in individuals over 80 years old, we investigated the relationship between age, antibody and Bmem levels.^89^ We found no correlation between RBD-specific IgG levels or RBD-specific Bmem numbers and age in the ChAdOx1 group, but did observe an antigen-specific correlation between lower WH1 RBD-specific IgG and Bmem numbers and older age after BNT162b2 vaccination. Others have also found differences in SARS-CoV-2-specific antibody quantities across age groups, and together this supports the implementation of separate vaccination schedules for more high-risk groups such as the elderly, to boost protective IgG and Bmem levels.^89–92^

The third COVID-19 vaccine dose has been shown to significantly boost NAb levels against Omicron, regardless of the primary vaccine formulation.^93^ It will be of interest to also examine RBD-specific Bmem recognition of VoC after booster doses, as these memory cells will likely be the source of these increased NAb titers. As circulating Bmem are more robust than NAb over time, this may better indicate whether booster doses are providing sustained immune protection against emerging VoC. Additionally, the capacity of RBD-specific Bmem to bind a range of Omicron sublineages after a bivalent variant-based vaccine should be investigated compared to a standard WH1-based booster. This will help to determine whether an Omicron-based booster vaccine provides sufficient protection against the currently circulating BA.5 and other emerging sublineages, or whether the vaccine must be updated with each new variant, as is currently done with seasonal quadrivalent influenza vaccines.^94^

Overall, we demonstrate that despite reduced antibody responses, ChAdOx1 vaccination induces robust circulating Bmem against SARS-CoV-2 Spike RBD with the capacity to recognize prevalent VoC. This understanding is critical when considering the global use of ChAdOx1 as a primary vaccination. As the virus and pandemic evolve, with the inevitable emergence of new VoC, a better understanding will be needed of the immunological mechanisms that underlie protection against severe disease. Considering the function of Bmem to readily respond upon renewed antigen encounter, detailed insights into their antigen specificity and affinity, as well as functional characteristic could provide guidance for the timing and need for future booster vaccinations.

## ACKNOWLEDGEMENTS

We thank the ARAFlowCore staff for training and assistance with flow cytometry, Ms Sandra Esparon, Ms Reema Bajaj, and Dr Bruce D Wines (Burnet Institute) for assistance with protein production, and Mr Jack Edwards, Ms Ebony Blight, and Ms Pei Mun Aui (Monash University) for sample collection and preparation. Supported by an ARA Honours Scholarship (HAF), Australian Government Research Training Program Scholarship (GEH), the Australian Government Medical Research Future Fund (MRFF, Project no. 2016108; MCvZ, HED and REO’H), the Victorian Operational Infrastructure Support Program (Burnet Institute), and an unrestricted research grant from BD Biosciences.

## AUTHOR CONTRIBUTIONS

Study design: HAF, GEH, ESJE, SJB, PMH, HED, REO’H and MCvZ; Performed experiments: HAF, GEH, NV, IB, and MCvZ; Formal analysis: HAF, GEH, NV, and IB; Provided reagents: SJB; Supervised the work: ESJE, REO’H and MCvZ; Wrote the manuscript: HAF and MCvZ. All authors edited and approved the final version of the manuscript.

