## Supplemental Material for "COVID-19 adenoviral vector vaccination elicits a robust memory B cell response with the capacity to recognize Omicron BA.2 and BA.5 variants"

### Supplementary Tables (n=3) and Figures (n=6)

### Supplementary Table 1. Participant characteristics

| Participant number | Age at inclusion (year) | Sex (M/F) | Time between doses of ChAdOx1 (days) |
|---|---|---|---|
| **1** | 26 | M | 84 |
| **2** | 27 | F | 84 |
| **3** | 29 | F | 84 |
| **4** | 32 | F | 78 |
| **5** | 33 | F | 83 |
| **6** | 33 | F | 84 |
| **7** | 33 | M | 84 |
| **8** | 34 | F | 82 |
| **9** | 35 | M | 84 |
| **10** | 36 | F | 84 |
| **11** | 37 | F | 83 |
| **12** | 37 | F | 84 |
| **13** | 38 | F | 84 |
| **14** | 42 | M | 84 |
| **15** | 43 | F | 84 |
| **16** | 45 | F | 84 |
| **17** | 45 | F | 84 |
| **18** | 47 | F | 76 |
| **19** | 48 | F | 88 |
| **20** | 48 | F | 83 |
| **21** | 48 | F | 85 |
| **22** | 50 | F | 79 |
| **23** | 54 | F | 84 |
| **24** | 56 | F | 91 |
| **25** | 57 | M | 84 |
| **26** | 58 | F | 84 |
| **27** | 58 | F | 95 |
| **28** | 59 | F | 70 |
| **29** | 61 | M | 84 |
| **30** | 64 | M | 84 |
| **31** | 65 | M | 84 |
| | Median: 45 (range: 26-65 years) | 74% female | Median: 84 (range: 70-95 days) |

### Supplementary Table 2. Flow cytometry antibody panel composition

| **Tube** | **Fluorochrome** | | | | | | | | | | | | | | | |
|---|---|---|---|---|---|---|---|---|---|---|---|---|---|---|---|---|
| | BUV395 | BUV496 | BUV737 | BV421 | BV480 | BV650 | BV711 | BV786 | FITC | BB700/ PerCP Cy-5.5 | PE | PE-Vio615 | PC7/ PE-Cy7 | APC | AF700 | APC-H7/ APC-Cy7 |
| **1. Trucount** | — | — | — | — | — | — | — | — | CD3 | CD45 | CD16+ CD56 | — | CD4 | CD19 | — | CD8a |
| **2. SARS-CoV-2-specific Bmem** | WH1 RBD | CD3 | WH1 RBD | CD27 | BA.2 RBD | BA.5 RBD | CD21 | CD71 | IgG2+ IgG3 | IgD | IgG1+ IgG2 | IgA | CD19 | IgG4 | Fixable viability | CD38 |
| **3. Streptavidin control** | Streptavidin | — | Streptavidin | CD27 | Streptavidin | Streptavidin | — | — | CD3 | IgD | — | — | CD19 | — | Fixable viability | — |

### Supplementary Table 3: Details of antibodies used in flow cytometry panels

| Marker | Fluorochrome | Clone | Vendor | Cat. number | Volume /test (µL) | Tube |
|---|---|---|---|---|---|---|
| CD3 | BUV496 | UCHT1 | BD Biosciences | 612940 | 1 | 2 |
| CD3 | FITC | UCHT1 | BD Biosciences | 555332 | 1 | 3 |
| CD3 | FITC | SK7 | BD Biosciences | 662995* | 46ng | 1 |
| CD4 | PE-Cy7 | SK3 | BD Biosciences | 662995* | 30ng | 1 |
| CD8a | APC-Cy7 | SK1 | BD Biosciences | 662995* | 126ng | 1 |
| CD16 | PE | B73.1 | BD Biosciences | 662995* | 33ng | 1 |
| CD19 | APC | SJ25C1 | BD Biosciences | 662995* | 46ng | 1 |
| CD19 | PE-Cy7 | SJ25C1 | BD Biosciences | 557835 | 5 | 2, 3 |
| CD21 | BV711 | B-ly4 | BD Biosciences | 563163 | 5 | 2 |
| CD27 | BV421 | M-T271 | BD Biosciences | 562513 | 1 | 2, 3 |
| CD38 | APC-H7 | HB7 | BD Biosciences | 656646 | 1 | 2 |
| CD45 | PerCP Cy-5.5 | 2D1 | BD Biosciences | 662995* | 120ng | 1 |
| CD56 | PE | NCAM16.2 | BD Biosciences | 662995* | 22ng | 1 |
| CD71 | BV786 | M-A712 | BD Biosciences | 563768 | 1 | 2 |
| Fixable viability | AF700 | N/A | BD Biosciences | 564997 | 0.1 | 2, 3 |
| IgA | PE-Vio615 | REA1014 | Miltenyi Biotec | 130-116-882 | 1.5 | 2 |
| IgD | BB700 | IA6-2 | BD Biosciences | 566538 | 1 | 2, 3 |
| IgG1 | PE | G17-1 | BD Biosciences | 624049 | 0.1 | 2 |
| IgG2 | FITC | HP6002 | BD Biosciences | 624045 | 0.5 | 2 |
| IgG2 | PE | HP6002 | BD Biosciences | 624049 | 1 | 2 |
| IgG3 | FITC | HP6047 | BD Biosciences | 624045 | 0.5 | 2 |
| IgG4 | APC | SAG4 | Cytognos | CYT-IGG4AP | 2 | 2 |
| Streptavidin | BUV395 | - | BD Biosciences | 564176 | 0.67 | 2, 3 |
| Streptavidin | BUV737 | - | BD Biosciences | 612775 | 0.67 | 2, 3 |
| Streptavidin | BV480 | - | BD Biosciences | 564876 | 0.67 | 2, 3 |
| Streptavidin | BV650 | - | BioLegend | 563855 | 0.13 | 2, 3 |

*Antibodies part of the Multitest$^{TM}$ 6-color TBNK kit (BD Biosciences)

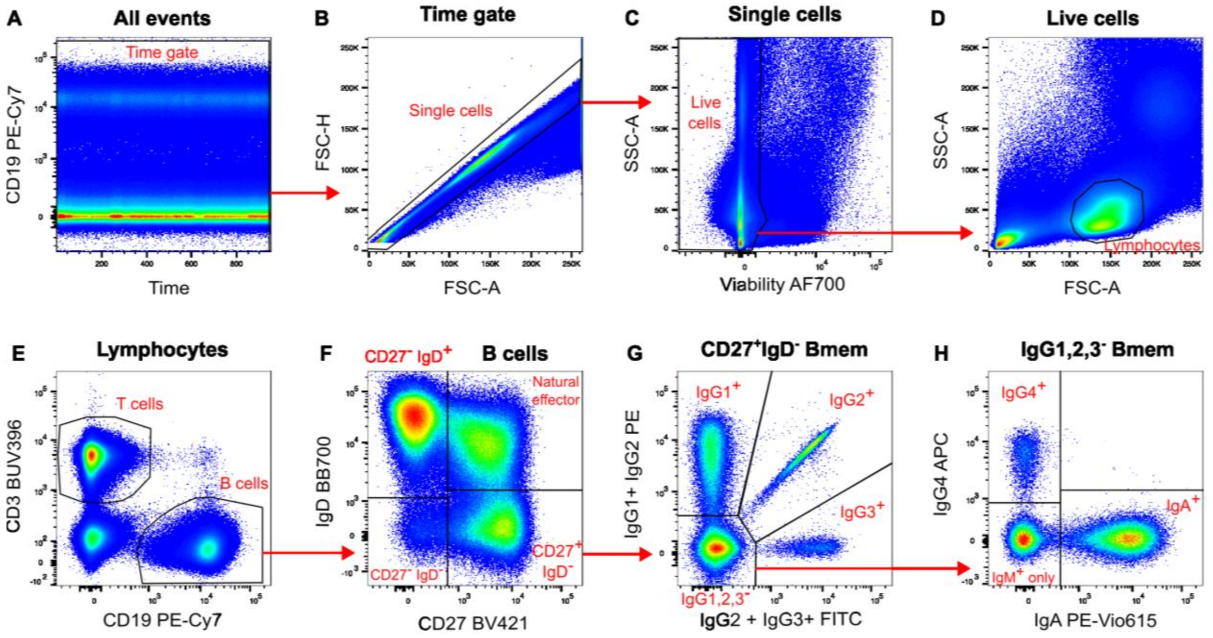


**Supplementary Figure 1. B-cell gating strategy.** (**A**) CD19 vs Time was used to ensure a steady signal of CD19 PE-Cy7. (**B**) Doublets were excluded by gating on FSC-A vs FSC-H. (**C**) Dead cells were excluded by gating on SSC-A vs Viability. (**D**) Live cells were gated for SSC$^{lo}$ lymphocytes. (**E**) Lymphocytes were gated for CD3$^+$CD19$^-$ T cells and CD3$^-$CD19$^+$ B cells. (**F**) B cells were gated for CD27$^-$IgD$^+$ naive B cells and CD27$^+$IgD$^+$, CD27$^+$IgD$^-$, and CD27$^-$IgD$^-$ Bmem subsets. (**G**) Each Bmem subset was gated for IgG1$^+$, IgG2$^+$, and IgG3$^+$ populations, then (**H**) the triple negative population was further divided into IgG4$^+$, IgA$^+$, and IgM$^+$ only cells. Representative plots from donor sample taken post-dose two of ChAdOx1.

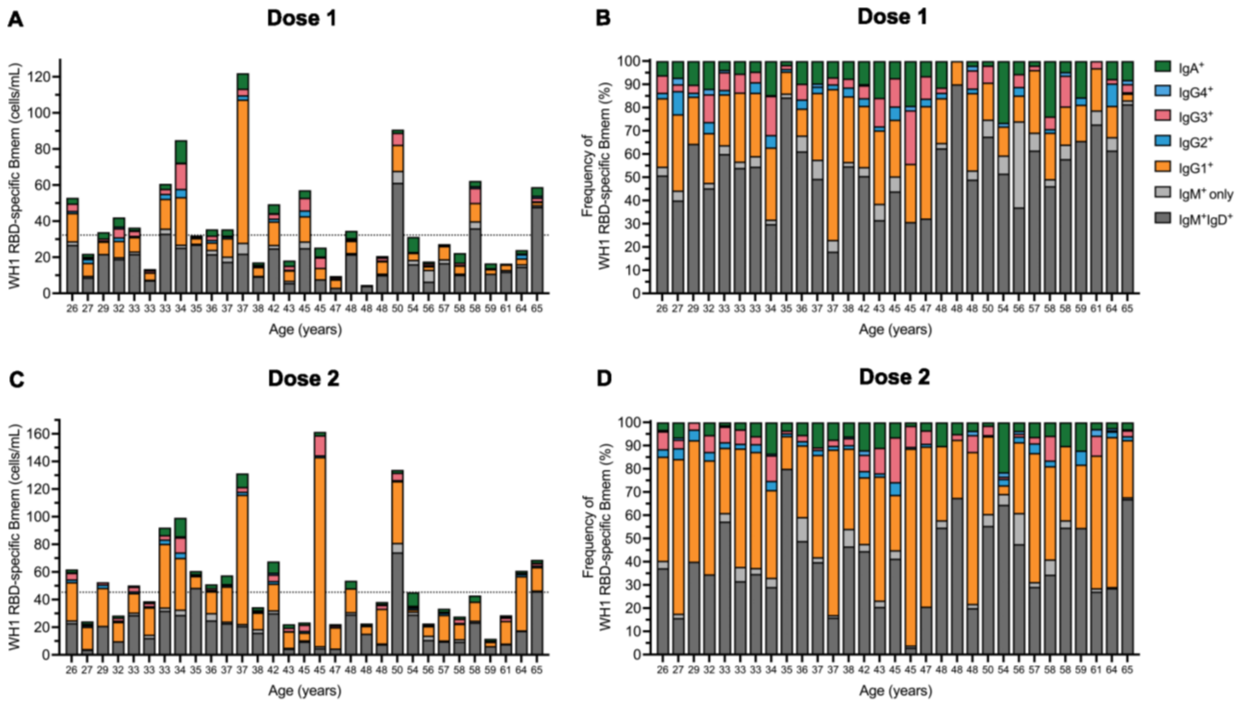


**Supplementary Figure 2. Ig isotype and IgG subclass distribution in individual donors.** (**A**) Absolute numbers and (**B**) relative distribution of WH1 RBD-specific Bmem expressing IgG1, IgG2, IgG3, IgG4, IgA, IgM only, or IgM and IgD post-dose one of ChAdOx1. (**C**) Absolute numbers and (**D**) relative distribution of each Ig isotype and IgG subclass post-dose two of ChAdOx1. Dotted lines represent median WH1 RBD-specific Bmem numbers.

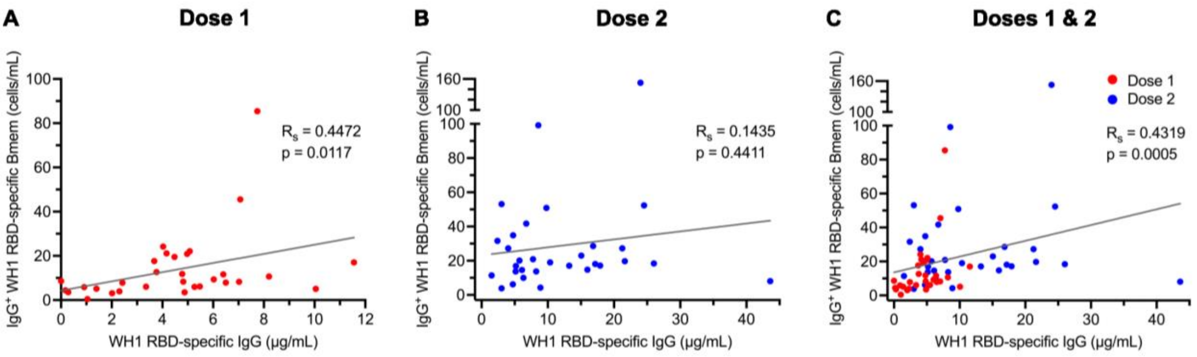


**Supplementary Figure 3. Correlation between IgG$^{+}$ WH1 RBD-specific Bmem and plasma IgG.** Correlation between absolute numbers of IgG$^{+}$ WH1 RBD-specific Bmem and WH1 RBD-specific plasma IgG (A) post-dose one, (B) post-dose two, and (C) combined post-dose one and two of ChAdOx1. Non-parametric Spearman's rank correlation ($R_s$), solid line represents simple linear regression line.

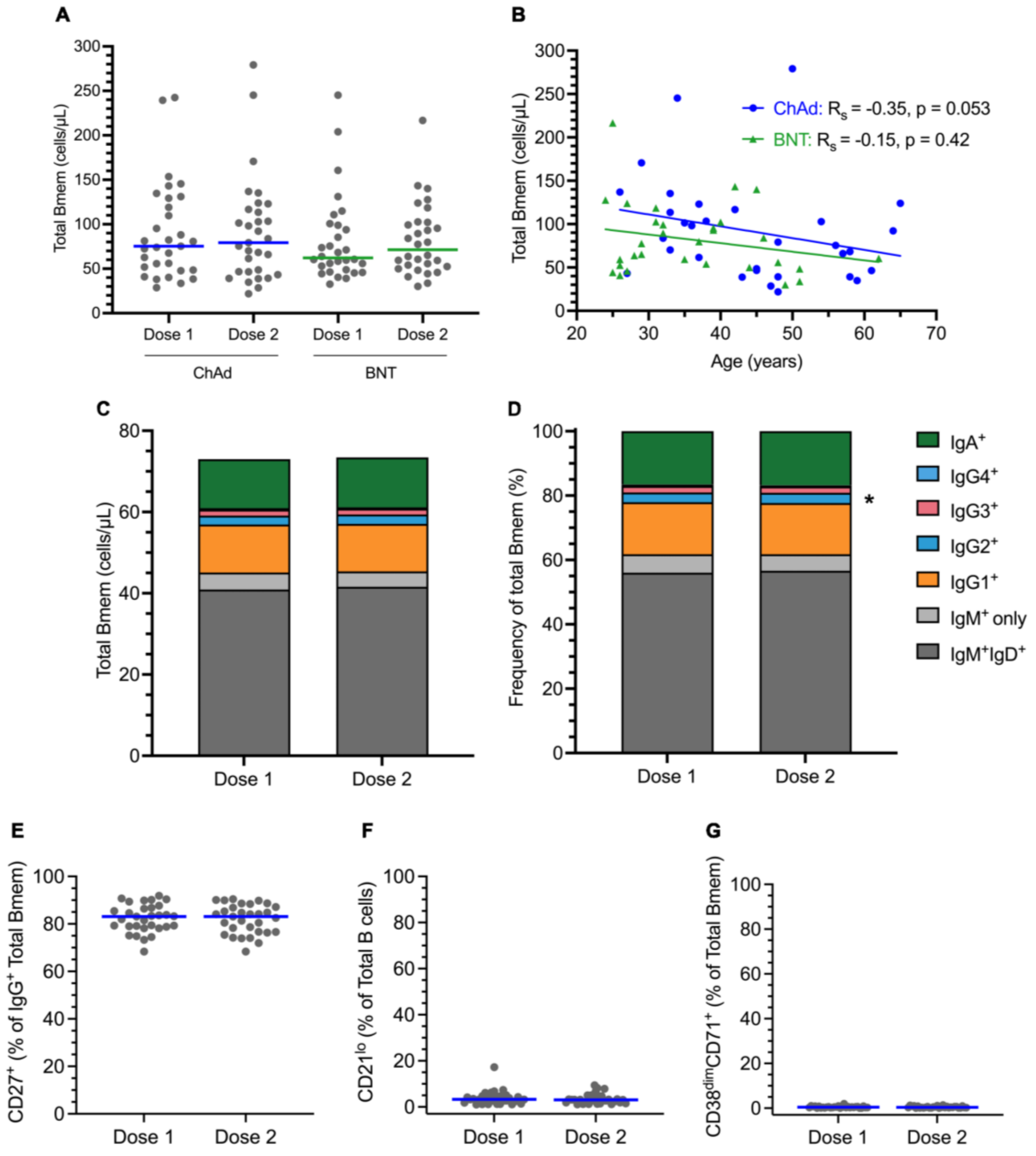


**Supplementary Figure 4. Immunophenotype of total Bmem following ChAdOx1 vaccination.** (**A**) Absolute numbers of total Bmem four weeks post-dose one (Dose 1) and four weeks post-dose two (Dose 2) of ChAdOx1 (ChAd) and BNT162b2 (BNT). (**B**) Correlation of total Bmem with age after either ChAdOx1 or BNT162b2 two-dose vaccination. Non-parametric Spearman's rank correlation ($R_S$), solid line represents simple linear regression line. (**C**) The percentage of total Bmem expressing IgM only, IgM and IgD, IgG1, IgG2, IgG3, IgG4, or IgA four weeks post-dose one (Dose 1) and four weeks post-dose two (Dose 2) of ChAdOx1. (**D**) The percentage of WH1 RBD-specific Bmem expressing each Ig isotype and IgG subclass postvaccination with ChAdOx1. (**E**) Frequencies of total $IgG^+$ Bmem expressing CD27 post-vaccination with ChAdOx1. (**F-G**) The frequencies of total (**F**) $CD21^{lo}$ and (**G**) $CD38^{dim}CD71^+$ Bmem post-vaccination with ChAdOx1. ChAdOx1 n=31, BNT162b2 n=30. Wilcoxon matched-pairs singed rank test. Only significant differences shown. *p<0.05, ****p<0.0001. BNT162b2 data previously published.[1]

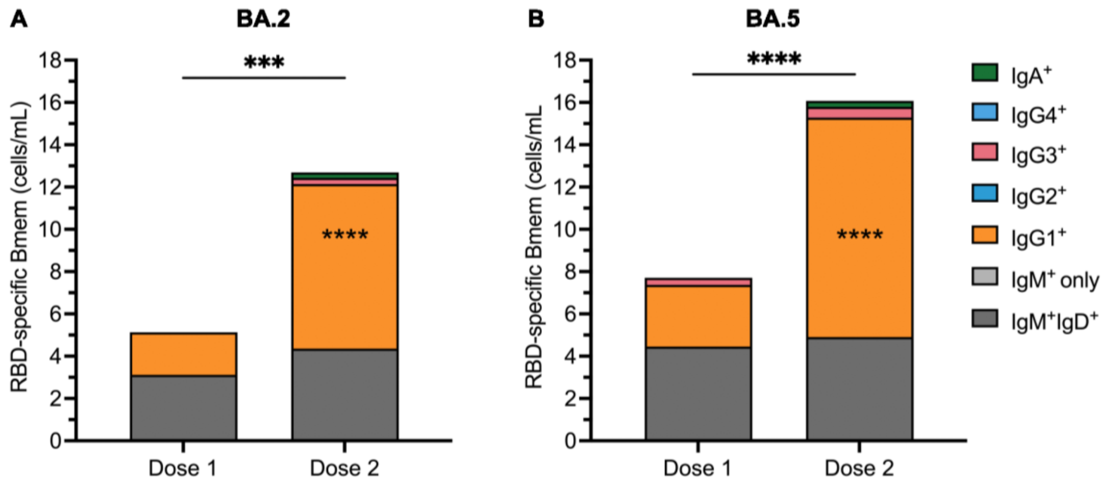


**Supplementary Figure 5. Relative distribution of the Ig isotypes and IgG subclasses of BA.2 and BA.5 RBD-specific Bmem following ChAdOx1 vaccination.** (**A**) The percentage of RBD-specific Bmem binding BA.2 expressing IgM only, IgM and IgD, IgG1, IgG2, IgG3, IgG4, or IgA four weeks post-dose one (Dose 1) and four weeks post-dose two (Dose 2) of ChAdOx1. (**B**) The percentage of RBD-specific Bmem binding BA.5 Bmem expressing each Ig isotype and IgG subclass postvaccination with ChAdOx1. n=31. Wilcoxon matched-pairs singed rank test. Only significant differences shown. ***p<0.001, ****p<0.0001.

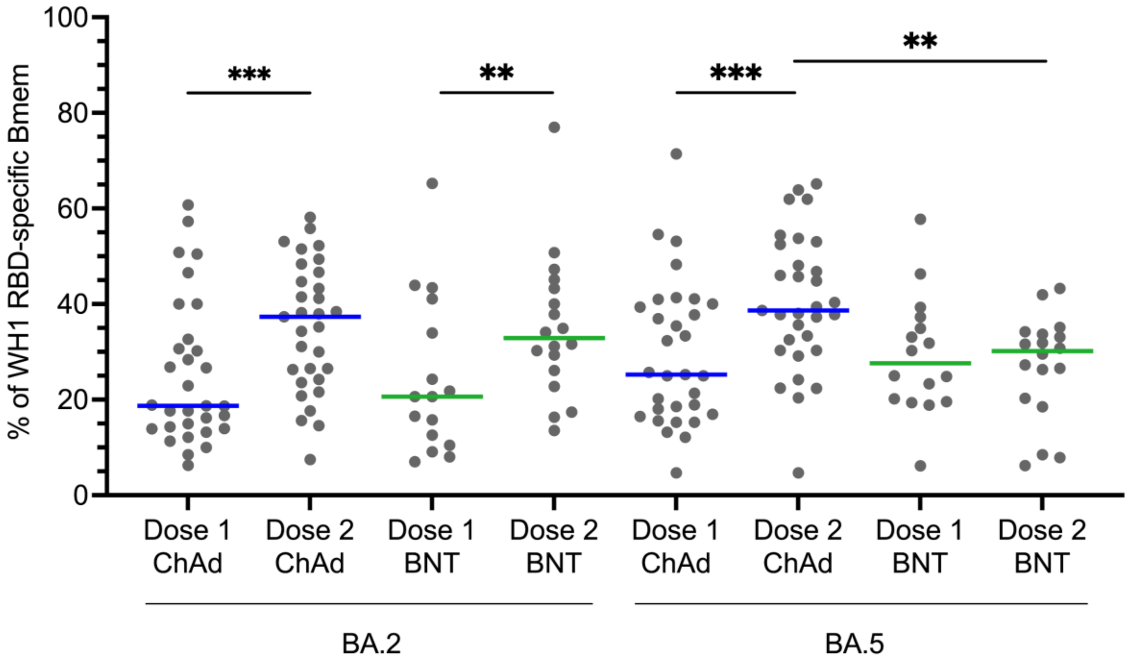


**Supplementary Figure 6. Comparison of the capacity of WH1 RBD-specific Bmem to bind Omicron BA.2 and BA.5 after vaccination with ChAdOx1 or BNT162b2.** Frequencies of WH1 RBD-specific Bmem that bound BA.2 and BA.5 RBD post-vaccination with ChAdOx1 (ChAd, n=31) or BNT162b2 (BNT, n=16 post-dose one and n=18 post-dose two). Solid lines indicate median values. Mann-Whitney test for unpaired data and Wilcoxon matched-pairs signed rank test for paired data. Only significant differences shown. **p<0.01, ***p<0.001. BNT162b2 data previously published.[1]
